# Spike activity regulates vesicle filling at a glutamatergic synapse

**DOI:** 10.1101/2019.12.20.873083

**Authors:** Dainan Li, Yun Zhu, Hai Huang

## Abstract

Synaptic vesicles need to be recycled and refilled rapidly to maintain high-frequency synaptic transmission. However, little is known about the control of transport of neurotransmitter into synaptic vesicles, which determines the contents of synaptic vesicles and the strength of synaptic transmission. Here we report that Na^+^ substantially accumulated in the calyx of Held terminals of mouse during high-frequency spiking. The activity-induced elevation of cytosolic Na^+^ activated vesicular Na^+^/H^+^ exchanger, facilitated glutamate loading into synaptic vesicles and increased quantal size of asynchronous released vesicles, but did not affect the vesicle pool size or release probability. Consequently, presynaptic Na^+^ increased the excitatory postsynaptic currents and was required to maintain the reliable high-frequency signal transmission from the presynaptic calyces to the postsynaptic MNTB neurons. Blocking Na^+^/H^+^ activity with EIPA decreased the postsynaptic current and caused failures in postsynaptic firing. Therefore, during high-frequency synaptic transmission, when large amounts of glutamate are released, Na^+^ accumulated in the terminals, activated vesicular Na^+^/H^+^ exchanger, and regulated glutamate loading as a function of the level of vesicle release.

**Significant statement:** Auditory information is encoded by action potentials phase-locked to sound frequency at high rates. Large amount of synaptic vesicles are released during high-frequency synaptic transmission, accordingly, synaptic vesicles need to be recycled and refilled rapidly. We have recently found that a Na^+^/H^+^ exchanger expressed on synaptic vesicles promotes vesicle filling with glutamate. Here we showed that during high-frequency signaling, when massive vesicles are released, Na^+^ accumulates in terminals and facilitates glutamate uptake into synaptic vesicle. Na^+^ thus accelerates vesicle replenishment and sustains reliable synaptic transmission.

## Introduction

High-frequency firing neurons are widely distributed throughout the central nervous system, including the auditory brainstem, cerebellum, thalamus, hippocampus, and neocortex (Chen and Regehr, 1999; Hu and Jonas, 2014; McCormick et al., 1985; Rudy and McBain, 2001; Taschenberger et al., 2002). Accordingly, synaptic vesicles need to be recycled and refilled rapidly to support the high-frequency synaptic signaling (Edwards, 2007; Farsi et al., 2017; Rizzoli and Betz, 2005). Accumulating studies have uncovered the mechanisms of vesicle fusion and recycling (Heuser and Reese, 1973; Klingauf et al., 1998; Sudhof, 2004; Wang and Kaczmarek, 1998); however, the control of the contents of synaptic vesicles has received considerably less attention (Balmer and Trussell, 2016). Recent studies showed that synaptic vesicles express Na^+^/H^+^ monovalent cation exchanger (NHE) activity that converts the pH gradient into an electrical potential required by the vesicular glutamate transporter (Goh et al., 2011; Preobraschenski et al., 2014). Na^+^ flux through HCN channels enhances presynaptic Na^+^ concentration and thus promotes synaptic vesicle filling with glutamate (Huang and Trussell, 2014). However, how spike activity controls the presynaptic Na^+^ dynamics and how accumulated Na^+^ modulates the synaptic transmission are unknown. Using the mouse calyx of Held, a giant glutamatergic synapse in the medial nucleus of the trapezoid body (MNTB) of the auditory brainstem that permits direct pre- and postsynaptic recordings and manipulation of the presynaptic cytosolic environment, we showed that glutamate loading is facilitated by intracellular Na^+^ over the physiological concentration range. During high-frequency signaling, when large amounts of glutamate are released, Na^+^ accumulates in terminals, activates NHE, facilitates glutamate uptake into synaptic vesicles, thus accelerating vesicle refilling and sustaining reliable synaptic transmission. Thus presynaptic cytosolic Na^+^ works as a signaling ion to coordinate glutamate loading as a function of the level of vesicle release.

## Materials and Methods

### Slice Preparation

All animal handling and procedures were approved by the Institutional Animal Care and Use Committee of Tulane University and followed U.S. Public Health Service guidelines. Coronal brainstem slices containing the medial nucleus of the trapezoid body (MNTB) were prepared from C57BL/6J mice of either sex aged postnatal day 8-12, being similar to previously described (Zhang and Huang, 2017). Briefly, mice brainstems were dissected and 210 μm sections were sliced using a vibratome (VT1200S, Leica) in ice-cold, low-Ca^2+^, low-Na^+^ saline contained (in mM) 230 sucrose, 25 glucose, 2.5 KCl, 3 MgCl_2_, 0.1 CaCl_2_, 1.25 NaH_2_PO_4_, 25 NaHCO_3_, 0.4 ascorbic acid, 3 *myo*-inositol, and 2 Na-pyruvate, bubbled with 95% O_2_/5% CO_2_. Slices were immediately incubated at 32°C for 20-40 min and subsequently stored at room temperature in normal artificial cerebrospinal fluid (aCSF) contained (in mM): 125 NaCl, 25 glucose, 2.5 KCl, 1.8 MgCl_2_, 1.2 CaCl_2_, 1.25 NaH_2_PO_4_, 25 NaHCO_3_, 0.4 ascorbic acid, 3 myo-inositol, and 2 Na-pyruvate, pH 7.4 bubbled with 95% O_2_/5% CO_2_.

### Whole-Cell Recordings

Slices were transferred to a recording chamber and perfused with normal aCSF (2-3 ml/min) warmed to ∼32 °C by an in-line heater (Warner Instruments). Neurons were visualized using an Olympus BX51 microscope with a 60× water-immersion objective and custom infrared Dodt gradient contrast optics. Whole-cell patch-clamp recordings were performed with a Multiclamp 700B amplifier (Molecular Devices) except for the capacitance measurements. Pipettes pulled from thick-walled borosilicate glass capillaries (WPI) had open tip resistances of 3–5 MΩ and 2– 3 MΩ for the pre- and postsynaptic recordings, respectively. Series resistance (4-15 MΩ) was compensated by up to 70% (bandwidth 3 kHz). To record presynaptic action potentials under different Na^+^ concentration, pipette solution contained (in mM): 60 K-methanesulfonate, 20 KCl, 10 HEPES, 0.5 EGTA, 4 Mg-ATP, 0.3 Tris_3_-GTP, 14 Tris_2_-phosphocreatine, 5 glutamate, as well as 40 NMDG-methanesulfonate (for Na^+^-free solution), 10 Na-methanesulfonate + 30 NMDG-methanesulfonate (for 10 mM Na^+^ solution), or 40 Na-methanesulfonate (for 40 mM Na^+^ solution). All solutions were adjusted to pH 7.3 with KOH (290 mOsm) and the final K^+^ for all solutions was around 92 mM. For presynaptic voltage-clamp recordings, pipette solution contained (in mM): 70 Cs-methanesulfonate, 20 CsCl, 10 HEPES, 0.5 EGTA, 4 Mg-ATP, 0.3 Tris_3_-GTP, 10 Tris_2_-phosphocreatine, 5 Glutamate, as well as 40 NMDG-methanesulfonate (for Na^+^-free solution), 10 Na-methanesulfonate + 30 NMDG-methanesulfonate (for 10 mM Na^+^ solution), or 40 Na-methanesulfonate (for 40 mM Na^+^ solution). All solutions were adjusted to pH 7.3 with CsOH (310-315 mOsm). To isolate presynaptic Ca^2+^ currents in response to voltage steps, TEA-Cl (10 mM), and 4-AP (2 mM) and tetrodotoxin (1 µM) were added to ACSF, substituting for NaCl with equal osmolarity. For EPSC recordings, postsynaptic pipette solution contained (in mM): 130 Cs-methanesulfonate, 10 CsCl, 10 HEPES, 5 EGTA, 4 Mg-ATP, 0.3 Tris_3_-GTP, 5 Na_2_-phosphocreatine, 2 QX-314-Cl (290 mOsm, pH 7.3 with CsOH). Recording aCSF contained 5 μM (R)-CPP, 50 μM picrotoxin, and 1 μM strychnine to block NMDA, GABA, and glycine receptors, respectively. 2 mM kynurenic acid and 100 μM cyclothiazide were applied to block AMPA receptor saturation and desensitization, respectively. For postsynaptic spiking recordings, pipette solution contained (in mM): 135 K-gluconate, 10 KCl, 10 HEPES, 0.2 EGTA, 4 Mg-ATP, 0.3 Tris_3_-GTP, 7 Na_2_-phosphocreatine (290 mOsm, pH 7.3 with KOH). To record the asynchronous release, extracellular Ca^2+^ was replaced by Sr^2+^ (8 mM). Presynaptic action potentials were evoked by afferent fiber stimulation with a bipolar stimulating electrode placed close to the midline of slices. Liquid junction potentials were measured for all solutions, and reported voltages are appropriately adjusted.

### Perforated Patch-clamp Recordings

Perforated whole-cell patch-clamp recordings were used during prolonged high-frequency postsynaptic recordings. The pipette solution was similar to that for conventional whole-cell postsynaptic spiking recordings except gramicidin (final concentration of 60 µg/ml) was added immediately before use. The tip of the recording pipette was first filled with gramicidin-free solution and then back-fill with gramicidin-containing solution. The degree of perforation was monitored after formation of gigaohm seal and recordings were started after overall access resistance dropped to below 30 MΩ.

### Two-photon Na^+^-imaging

A Galvo multiphoton microscopy system (Scientifica) with a Ti:sapphire pulsed laser (Chameleon Ultra II; Coherent) was used for two-photon Na^+^ imaging (Bender et al., 2010). The laser was tuned to 810 nm, epifluorescence signals were captured through 60X, 1.0 NA objectives and a 1.4 NA oil immersion condenser (Olympus). Fluorescence was split into red and green channels using dichroic mirrors and band-pass filters. Data were collected in frame-scan or line-scan modes. Pipette solution contained (in mM): 110 K-methanesulfonate, 20 KCl, 10 HEPES, 0.5 EGTA, 4 Mg-ATP, 0.3 Tris_3_-GTP, 5 Na_2_-phosphocreatine (290 mOsm; pH 7.3 with KOH), while 1 mM SBFI and 20 μM Alexa 594 were added to the pipette solution before the experiments. Presynaptic spikes were evoked by afferent fiber stimulation and corresponding Na^+^ signals were recorded under two-photon imaging. Standard calibration methods were used to measure the absolute Na^+^ concentrations (Huang and Trussell, 2014; Rose, 2012).

### Membrane capacitance measurement

Whole-cell patch-clamp recordings were made with an EPC-10 USB double amplifier controlled by Patchmaster software (HEKA) at 32 °C to record presynaptic calcium currents and membrane capacitance (C_m_) responses. The calyx of Held terminals were voltage-clamped at holding potential of −80 mV with a sinusoidal wave (60 mV peak-to-peak amplitude at 1 kHz) superimposed (Sun and Wu, 2001). Tips of patch pipettes were coated with dental wax to reduce stray capacitance. Series resistance (6-20 MΩ) was electronically compensated by up to 75% (10 μs lag). The calyceal terminals were depolarized from –80 mV to +10 mV for 1 ms and 30 ms to mimic the action potential-induced release and to deplete the releasable pool, respectively. Recordings were performed. Data were obtained within 20 min after break-in and sampled at 100 kHz.

### Drugs

Drugs were obtained from Tocris (H-89), Alomone (tetrodotoxin, CPP, cyclothiazide), Invitrogen (SBFI and Alexa 594), and all others from Sigma-Aldrich.

### Analysis

Data were analyzed using Clampfit (Molecular Devices), Igor (WaveMetrics) and Image J. Data are expressed as mean ± S.E.M. Statistical significance was established using paired t-tests and one-way ANOVA as noted, with p < 0.05 indicating a significant difference.

## Results

### Spikes control presynaptic cytosolic Na+ concentration

Na^+^ flows into the cytosol during the firing of action potentials, while the location of Na^+^ channels in the calyx of Held is under debate (Huang and Trussell, 2008; Kim et al., 2010; Leao et al., 2005; Sierksma and Borst, 2017). Whole-cell recordings were made at the mouse calyx of Held and the presynaptic Na^+^ changes during action potential (AP) firing were assayed using two-photon laser scanning microscopy (Fig. 1). Na^+^ indicator SBFI (1 mM) and the volume marker Alexa 594 (15 μM) were loaded via patch pipettes. Calyces with complete heminode were used to monitor the Na^+^ changes at different locations under line-scans and frame-scans (Fig. 1A). The presynaptic [Na^+^] in the resting state was 15.8 ± 2.1 mM (n = 4). Upon 10 s stimulation at 100 Hz, Na^+^ accumulated into the whole calyx terminal (Fig. 1B-C). The Na^+^ rise and decay kinetics at axon heminode and calyceal terminal overlapped, while Na^+^ increase at the calyceal terminal (37.4 ± 4.8 mM) was slightly smaller than that of the axon heminode (42.7 ± 4.0 mM) (P = 0.009, n = 6). These results showed that spike activity substantially increase the cytosolic Na^+^ concentration in both axon heminode and presynaptic terminal.

**Figure 1.**
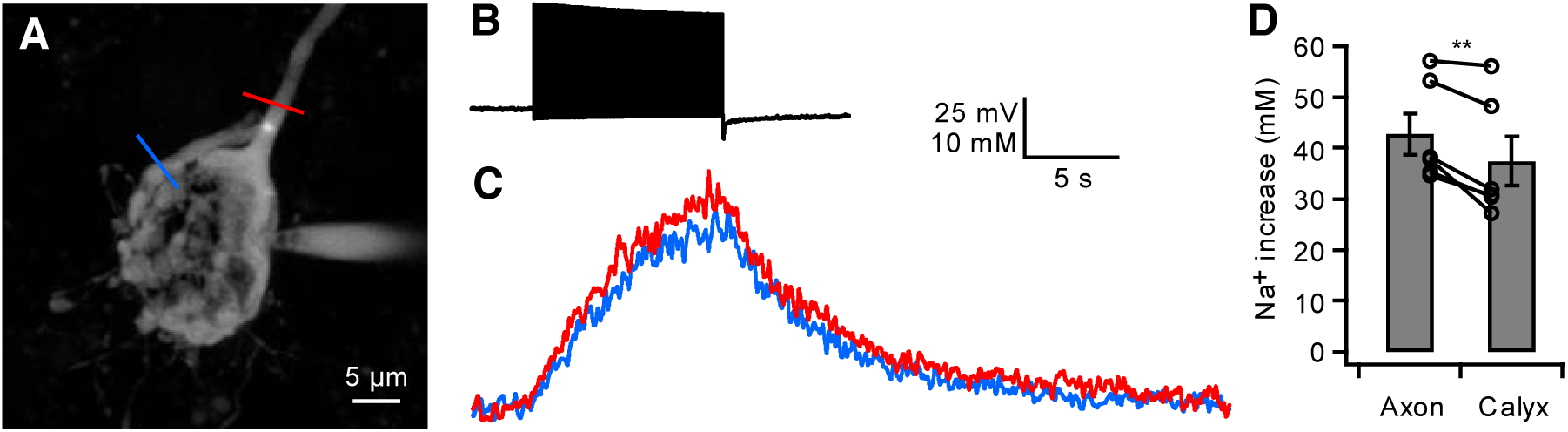
Presynaptic spikes control Na^+^ concentration. (A) Maximum intensity montage of a calyx of Held recorded with SBFI and Alexa 594 using two-photon microscopy. (B) Spikes were evoked at 100 Hz for 10 s by afferent fiber stimulation. (C) Corresponding Na^+^ increase were detected at both the preterminal axon heminode (red) and calyceal terminal (blue). (D) Summary plot of cytosolic Na^+^ increase at the axon and terminal.

### Presynaptic Na^+^; regulates glutamatergic transmission

Previous study showed that presynaptic cytosolic Na^+^ promotes vesicular glutamate uptake and miniature excitatory postsynaptic current (mEPSC) amplitude (Huang and Trussell, 2014). Here we performed simultaneous presynaptic and postsynaptic whole-cell recordings to examine how presynaptic Na^+^ influences action potential-driven synaptic transmission. We dialyzed the cytosolic contents of presynaptic calyces and simultaneously measured AMPA receptor-mediated excitatory postsynaptic currents (EPSCs) in MNTB principal neurons in the whole-cell mode. The calyces were recorded with pipette solutions containing 0 mM, 10 mM, or 40 mM Na^+^, pairs of presynaptic APs (10 ms intervals) were evoked every 15-20 s by afferent fiber stimulation and the resulting EPSCs were continuously recorded (Fig. 2). With the 10 mM Na^+^ solution presynaptically, the EPSC amplitude remained unchanged over 20 min of recording (104 ± 2%; *P* = 0.16, *n* = 6). When the calyceal terminals were broken into using a Na^+^-free pipette solution, the EPSC amplitude gradually declined by 23 ± 4% over the 20 min period (*P* = 0.003, *n* = 5), whereas increasing the presynaptic [Na^+^] to 40 mM induced a 39 ± 8% (*P* = 0.002, *n* = 8) increase in the EPSC amplitude (Fig. 2B-C). When normalized to the values observed immediately after break-in, the EPSC amplitude was clearly smaller in the Na^+^-free solution than in 10 mM Na^+^ (*P* = 0.0001) and higher in 40 mM Na^+^ compared to 10 mM Na^+^ (*P* = 0.003, unpaired t-test; Fig. 2C). The presynaptic action potentials remained stable over the recording period in all conditions (Fig. 2A). Since the change in Na^+^ is restricted to the nerve terminal and does not affect the postsynaptic neuron, these data indicate that the changes in EPSC amplitude reflected an alteration in presynaptic glutamate release. No apparent change of paired-pulse ratio (Fig. 2D) were observed in any of the groups, suggesting that the change in EPSC amplitude is unlikely to be caused by a change in the presynaptic release probability.

**Figure 2.**
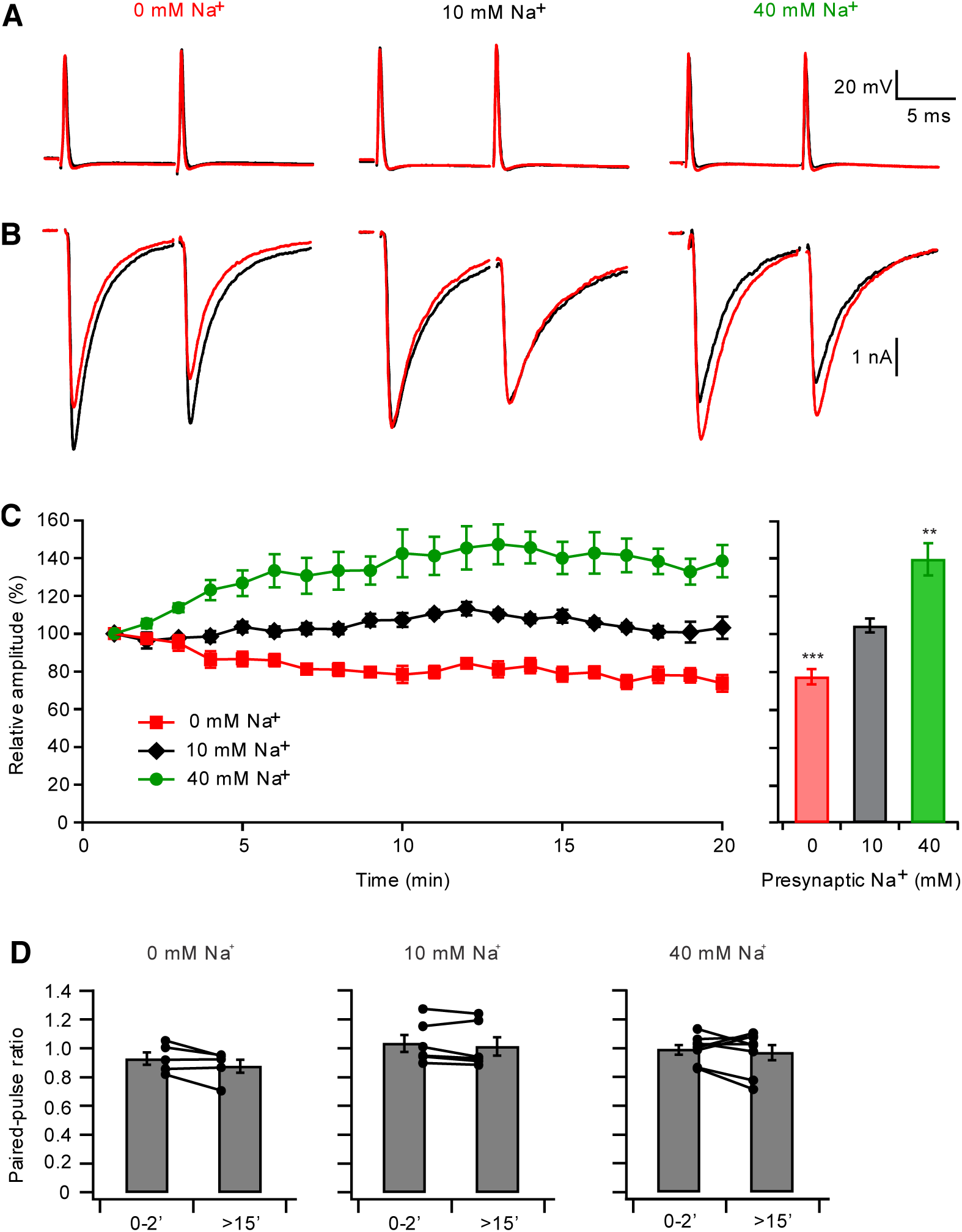
Presynaptic Na^+^ regulates the EPSC amplitude. (A) Pairs of presynaptic action potentials induced by afferent fiber stimulation when 0 mM (left), 10 mM (middle), or 40 mM Na^+^ (right) was present in the presynaptic pipette solution. Traces recorded within 2 min of break-in (black) and after 15 min (red) of recording are superimposed. (B) Postsynaptic currents in response to the presynaptic action potentials. (C) Left: time course of the normalized amplitudes of the first EPSC during paired-pulse recordings. Each point represents the average of 1 min recordings. Right: Relative amplitudes of the first EPSC after 15 min dialysis of the calyces with different Na^+^ concentrations. Amplitudes were normalized to the amplitudes measured within 2 min of break-in. (D) Presynaptic Na^+^ does not affect the paired-pulse ratio. Bar graphs of paired-pulse ratios measured within 2 min of break-in and after 15 min of recording were compared with presynaptic [Na^+^] of 0 mM (*P* = 0.12), 10 mM (*P* = 0.20), and 40 mM (*P* = 0.49). Statistical significance was assessed using a two-tailed, paired Student t-test. \*\**P* < 0.01. Error bars, ± S.E.M.

### Na^+^-dependent regulation of synaptic transmission is not due to change of AP waveform

Intracellular Na^+^ concentration will influence the Na^+^ driving force, affecting AP waveform, the Ca^2+^ influx and release of synaptic vesicles. We then used 1 ms depolarizing pulses from –80 mV to +10 mV to mimic the presynaptic action potential and trigger glutamate release (Fig. 3). Ca^2+^ currents remained stable over the recording period in all conditions (Fig. 3A). While the EPSC amplitude remained relative stable over 20 min of recording when the presynaptic solution contained 10 mM Na^+^ (95 ± 2%; *P* = 0.06, *n* = 5), the EPSC amplitude gradually declined by 20 ± 3% (*P* = 0.002, *n* = 5) with the Na^+^-free solution and increased by 32 ± 4% with 40 mM Na^+^ (*P* = 0.0002, *n* = 5; Fig. 3B-C). No change in the paired-pulse ratio (Fig. 3D) was observed in any group. This result was similar to that of the AP-triggered release, confirming that the presynaptic Na^+^-induced change in EPSC amplitude is not due to changes in AP waveform and Ca^2+^ influx.

**Figure 3.**
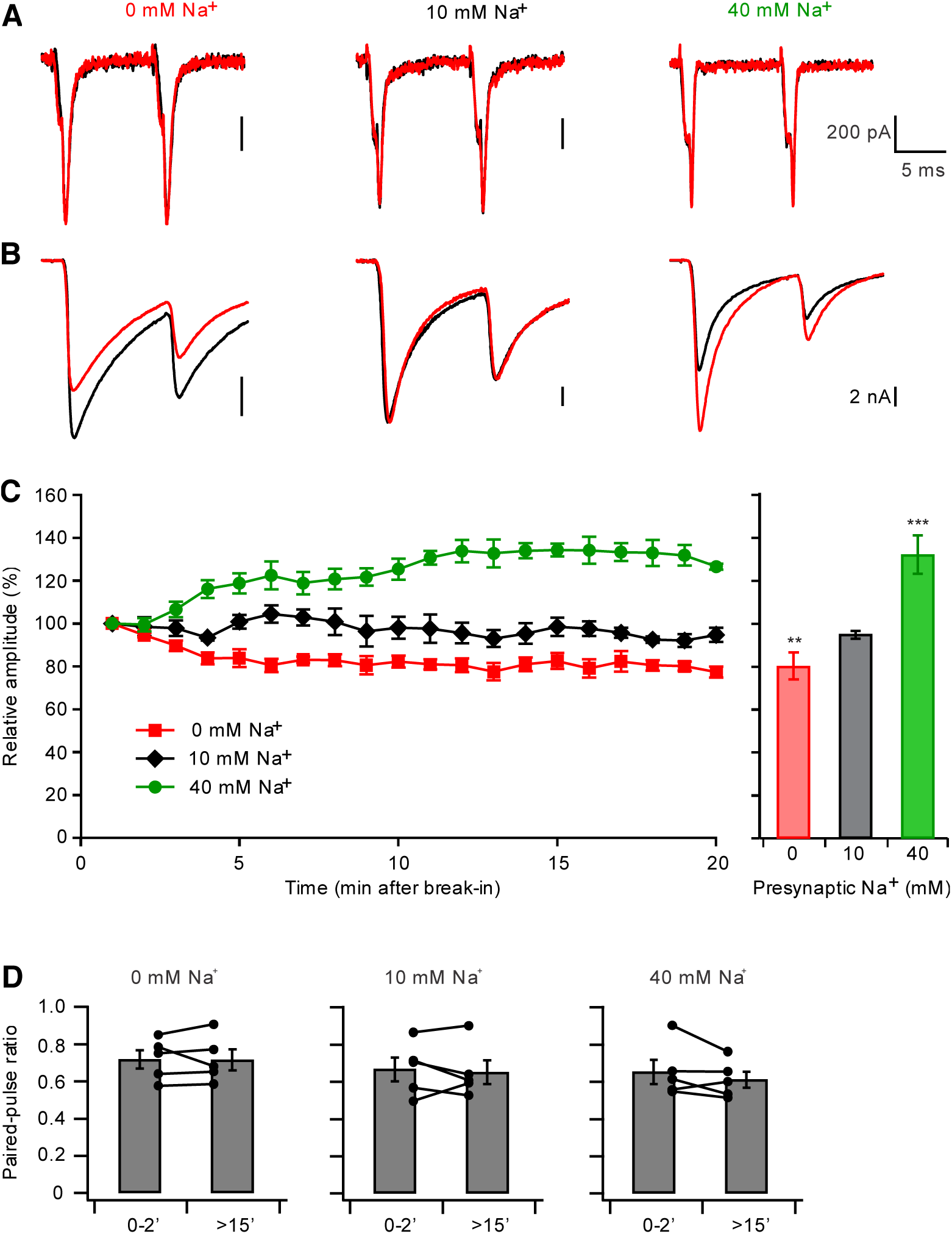
Presynaptic Na^+^ regulates EPSC amplitude in response to presynaptic depolarization. (A-B) Presynaptic calcium currents (A) and corresponding postsynaptic responses (B) induced by pairs of 1 ms presynaptic depolarizations from –80 mV to +10 mV when 0 mM (left), 10 mM (middle), or 40 mM (right) Na^+^ was present in the presynaptic pipette solutions. Recordings made within 2 min of break-in (black) and after 15 min (red) are superimposed. (C) Left: time course of the changes in EPSC amplitude. Each point represents the average of 1 min recordings. Right: Relative amplitudes of the first EPSC at 15-20 min of recordings with different presynaptic Na^+^ concentrations. Amplitudes were normalized to the amplitudes measured within 2 min of break-in. (D) Presynaptic Na^+^ does not affect the paired-pulse ratio in response to presynaptic depolarization. Bar graphs of paired-pulse ratios measured within 2 min of break-in and after 15 min of recording were compared with presynaptic [Na^+^] of 0 mM (p = 0.95), 10 mM (p = 0.69), and 40 mM (p = 0.25). Statistical significance was assessed using a two-tailed, paired Student t-test. \*\**P* < 0.01, ***p < 0.001. Error bars, ± S.E.M.

### Presynaptic Na^+^does not affect the synaptic vesicle pool or release probability

We next studied the mechanisms of presynaptic Na^+^ regulation of glutamatergic synaptic transmission. Measurement of membrane capacitance (C_m_) allows direct detection of exocytosis of synaptic vesicles at the calyx of Held with high temporal resolution (Sun and Wu, 2001). Typically, a 1-ms depolarization from –80 mV to +10 mV can elicit a Ca^2+^ influx and capacitance change (ΔC_m_) equivalent to a single action potential, while a 30-ms step depolarization is sufficient to deplete the whole readily releasable pool at calyceal terminals (Fedchyshyn and Wang, 2005; Renden and von Gersdorff, 2007; Wu and Borst, 1999). We found the ΔC_m_ evoked by a 1-ms step depolarization or by a 30-ms step depolarization showed no difference across these groups (*P* = 0.63 and 0.81, respectively, ANOVA test; Fig. 4). Release probability, defined as the ratio of single action potential-evoked ΔCm by the readily releasable pool, was not different among the different [Na^+^] groups (*P* = 0.39). These data suggest that the presynaptic [Na^+^] does not affect the presynaptic readily releasable pool size or the release probability.

**Figure 4.**
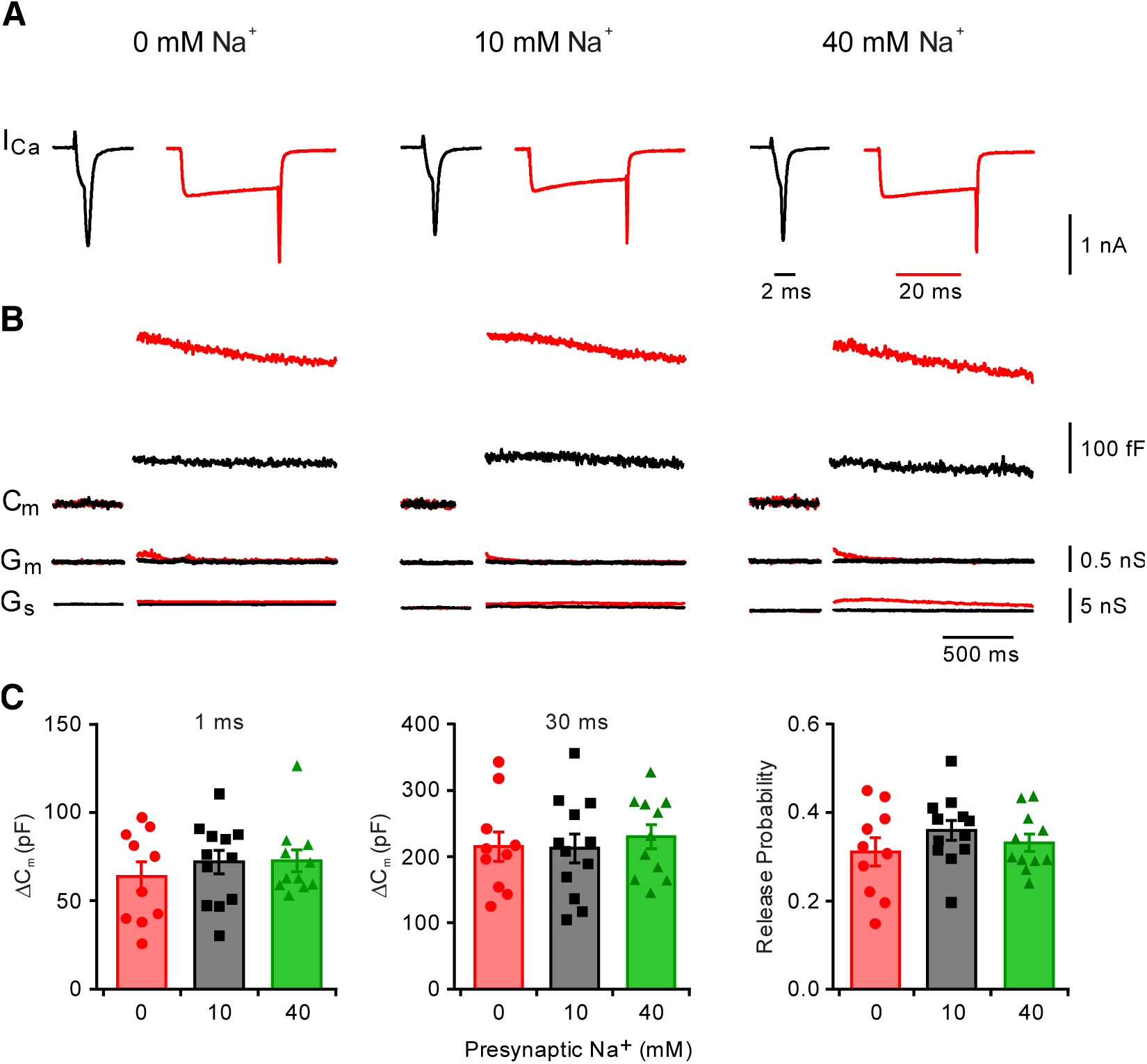
Presynaptic Na^+^ does not affect the readily releasable pool size or release probability. (a-b) Sampled presynaptic Ca^2+^ currents (I_Ca_) (A) and C_m_ responses (B) induced by a 1-ms (black) or 30-ms (red) depolarizations from –80 mV to +10 mV with presynaptic pipette solutions containing 0 mM (left), 10 mM (middle), or 40 mM (right) Na^+^. The corresponding membrane conductance (G_m_) and series conductance (G_s_) are shown to confirm the recording stability. (C) Group data show that the presynaptic [Na^+^] does not affect the C_m_ responses or release probability. Error bars, ±SEM.

### Presynaptic Na^+^ controls vesicular glutamate content

We next examined whether presynaptic Na^+^ regulates vesicular glutamate content. Recent studies showed that synaptic vesicles undergoing spontaneous and evoked fusion may derive from different pools (Chanaday and Kavalali, 2018; Fredj and Burrone, 2009). We recorded the amplitude of asynchronous EPSCs (aEPSCs) in response to single action potentials, since evoked synchronous release and asynchronous release share the same set of vesicles (Kaeser and Regehr, 2014). Single spike-triggered aEPSCs were induced by substituting extracellular Ca^2+^ with Sr^2+^ and were recorded in the postsynaptic MNTB principle neurons (Fig. 5). Immediately after presynaptic break-in to whole-cell mode (within 2 min), the aEPSCs showed no significant difference in amplitude in any of the groups (*P* = 0.20, one-way ANOVA test). With a 10 mM presynaptic Na^+^ solution, the aEPSC amplitude remained stable for over 10 min (98 ± 2%; *P* = 0.50, *n* = 5). The aEPSC amplitude was significantly reduced by 15 ± 2% (P = 0.003; n = 5) with 0 mM Na^+^ and increased by 17 ± 2% (*P* = 0.0005; n = 6) with 40 mM Na^+^ presynaptically. Bath application of 8-Br-cAMP, a cell membrane-permeant cAMP analog that activates HCN channels and increases presynaptic Na^+^ levels (Huang and Trussell, 2014), also increased the aEPSC amplitude by 19 ± 2% (*P* = 0.0006; n = 6). The difference in aEPSC amplitude suggested that alteration of the presynaptic Na^+^ level affected vesicular glutamate loading for both spontaneous and evoked release.

**Figure 5.**
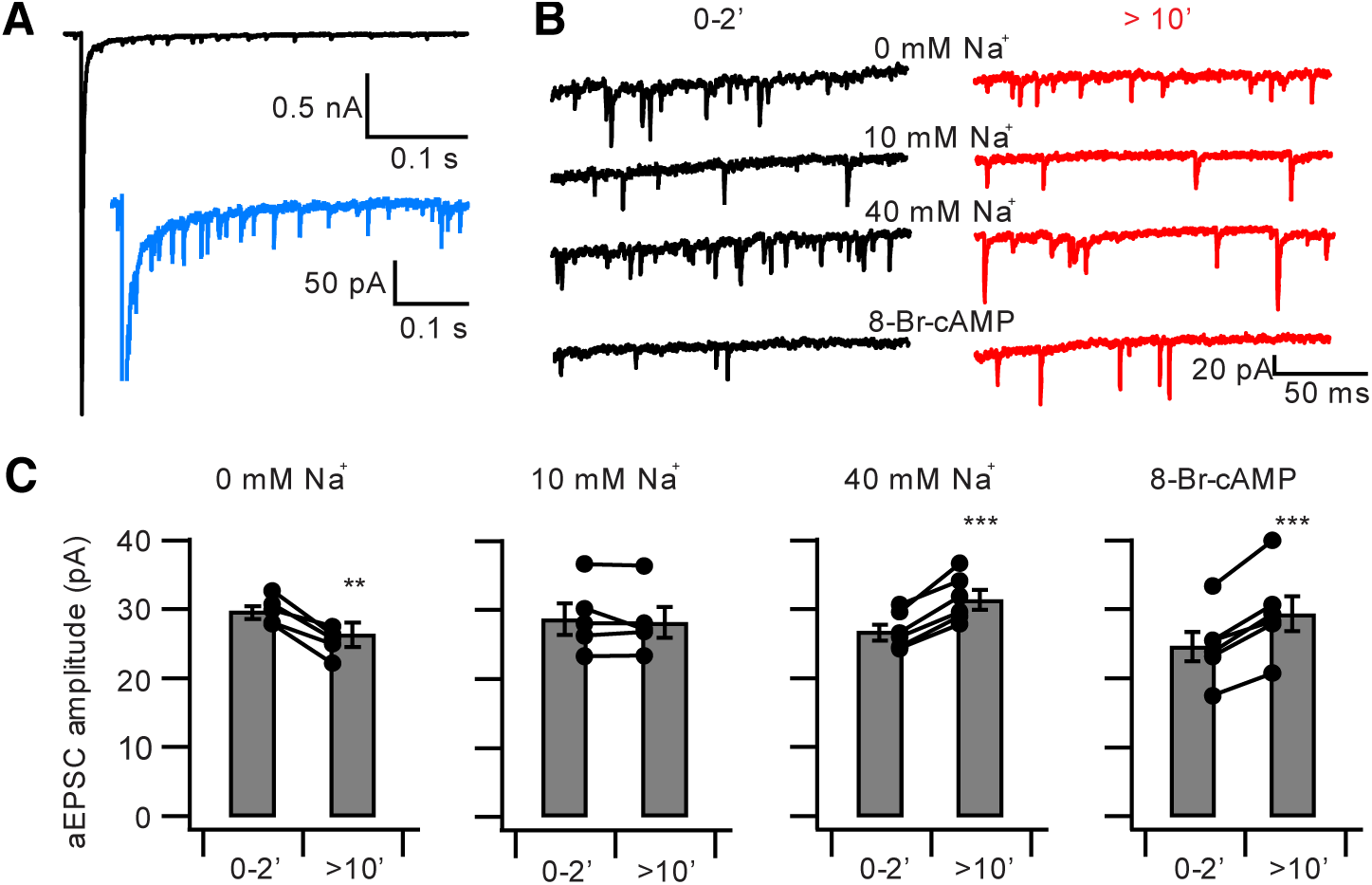
Presynaptic Na^+^ regulates aEPSC amplitude. (A) Example trace of prolonged period of aEPSCs following an initial evoked EPSC in response to a single stimulation. The insert shows an expanded trace of the asynchronous release (blue). (B) Left, example traces of asynchronous events within 2 min (black) or > 10 min (red) after presynaptic break-in with a 0 mM, 10 mM, or 40 mM Na^+^ patch pipette solution. The bottom trace shows the recordings before (control, black) and 10 min after application of 8-Br-cAMP (red); 10 µM H-89 was present in the postsynaptic pipette solution to inhibit 8-Br-cAMP induced kinase activation. (C) Bar graphs of the change in aEPSC amplitude in the 0 mM (n = 5, p = 0.003), 10 mM (n = 5, p = 0.50), 40 mM Na^+^ (n = 6, p = 0.0005), or 8-Br-cAMP (*n* = 6, *P* = 0.0006) groups. Statistical significance was assessed using a two-tailed paired Student t test. **p < 0.01; ***p < 0.001. Error bars, ± S.E.M.

### Presynaptic Na^+^ is required for reliable signal transmission

High-frequency signals of each globular bushy cell reliably transmit to a target MNTB principle neuron through the calyx of Held terminal, with few or no failures (Lorteije et al., 2009; Mc Laughlin et al., 2008). We next asked whether the cytosolic Na^+^-dependent modulation of EPSCs affects the reliability of signal transmission from the presynaptic calyx of Held to the postsynaptic MNTB neuron. Paired pre- and postsynaptic action potentials were recorded in response to calyceal fiber stimulation. Fiber stimuli at 200 Hz reliably evoked action potentials in the presynaptic terminals during the whole-cell recordings in 0, 10, and 40 mM Na^+^ conditions (Fig. 6A). Immediately after presynaptic break-in to whole-cell mode, most of the triggered presynaptic spikes correlated with postsynaptic spikes in the MNTB neurons, with only a few failures in the late phase of the stimulation (Fig. 6B). When the calyceal terminals were dialyzed with a 10 mM Na^+^ solution, no significant difference in the reliability of postsynaptic action potentials was observed over recording durations of >10 min (*P* = 0.46, n = 8, Fig. 6B-E). The probability of failure increased gradually with time when the presynaptic solution was Na^+^-free. After 10 min of presynaptic dialysis, the reliability was greatly reduced, and presynaptic release in the late part of the stimulus train was unable to drive postsynaptic action potentials (P = 0.01, n = 6, Fig. 6B-E). In contrast, when the calyceal terminal was dialyzed with a 40 mM Na^+^ solution, presynaptic spikes transmitted to the postsynaptic neuron showed less failures and higher firing probability after repeated stimuli trains, indicating that higher presynaptic Na^+^ level enhanced reliable synaptic transmission at the calyx of Held synapse (P = 0.0039, n = 8, Fig. 6B-E). It was notable that, for those events where the MNTB principle neuron failed to fire an action potential, a subthreshold EPSP was observed, indicating that the presynaptic spikes caused reliable release, but the synaptic current was not big enough to trigger a postsynaptic spike. Thus, the presynaptic Na^+^ level appears to play a regulatory role in facilitating vesicular glutamate uptake and eventually boosting reliable synaptic transmission.

**Figure 6.**
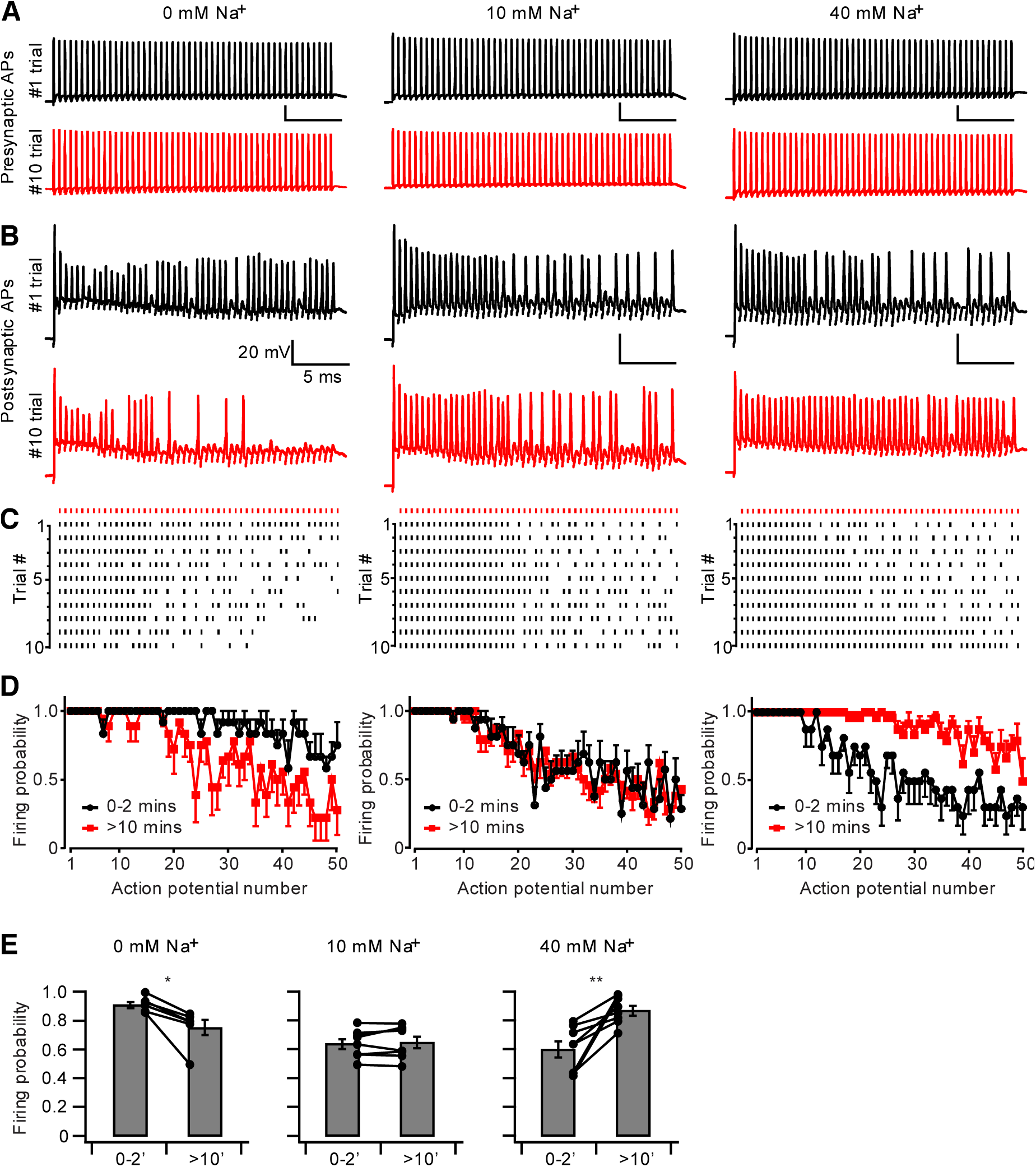
Presynaptic Na^+^ level contributes to reliable synaptic transmission. (A) Example traces of 50 presynaptic action potentials immediately (black) and 10 min (red) after break-in with 0 mM, 10 mM, and 40 mM Na^+^ in the presynaptic pipette solution. Action potentials were evoked by 200 Hz afferent fiber stimulation. (B) Postsynaptic spiking in MNTB principal neurons in response to the presynaptic firing in A. (C) Raster plots of spikes evoked by 200 Hz, 250 ms stimulus trains repeated with 60 s intervals. Presynaptic spikes are shown at the top in red. (D) Summary plots of average postsynaptic firing probability in response to 50 presynaptic action potentials at 200 Hz. (E) Overall change in the postsynaptic firing probability between the first 2 min and after 10 min, with 0 mM, 10 mM, or 40 mM presynaptic Na^+^. Statistical significance was assessed using a two-tailed paired Student t test. *p < 0.05, **p < 0.01; Error bars, ± SEM.

### NHE activity promotes synaptic transmission and signaling reliability

Synaptic vesicles express Na^+^/H^+^ monovalent cation exchanger (NHE) that converts the pH gradient into an electrical potential required by the vesicular glutamate transporter and promotes synaptic vesicle filling with glutamate (Goh et al., 2011; Huang and Trussell, 2014; Preobraschenski et al., 2014). Since presynaptic spikes substantially increase the presynaptic Na^+^ level, we asked if NHE activity is required in maintaining the high reliable synaptic transmission. Under perforated patch-clamp recordings, we were able to record EPSC and postsynaptic action potentials over a long duration (>30 min). Prolonged presynaptic fiber stimulation evoked reliable and stable EPSCs. Incubating EIPA (100 µM), an NHE specific blocker, reduced the EPSC amplitude to 54.8 ± 6.8% (p = 0.003, n = 5, Fig. 7A, C). Meanwhile, the postsynaptic firing started to fair after 6 min incubation of EIPA; the overall firing probability reduced to 21.2 ± 8.6% (P = 0.003, n = 4, Figure. 7B, D). A subthreshold EPSP was always observed when MNTB neuron failed to fire action potential, indicating the presynaptic spikes invaded into the terminals while the glutamate contents were not enough to trigger postsynaptic spike. Decrease of both EPSC amplitude and firing probability confirms Na^+^ and NHE activity is required in maintaining reliable synaptic signaling at high-frequency.

**Figure 7.**
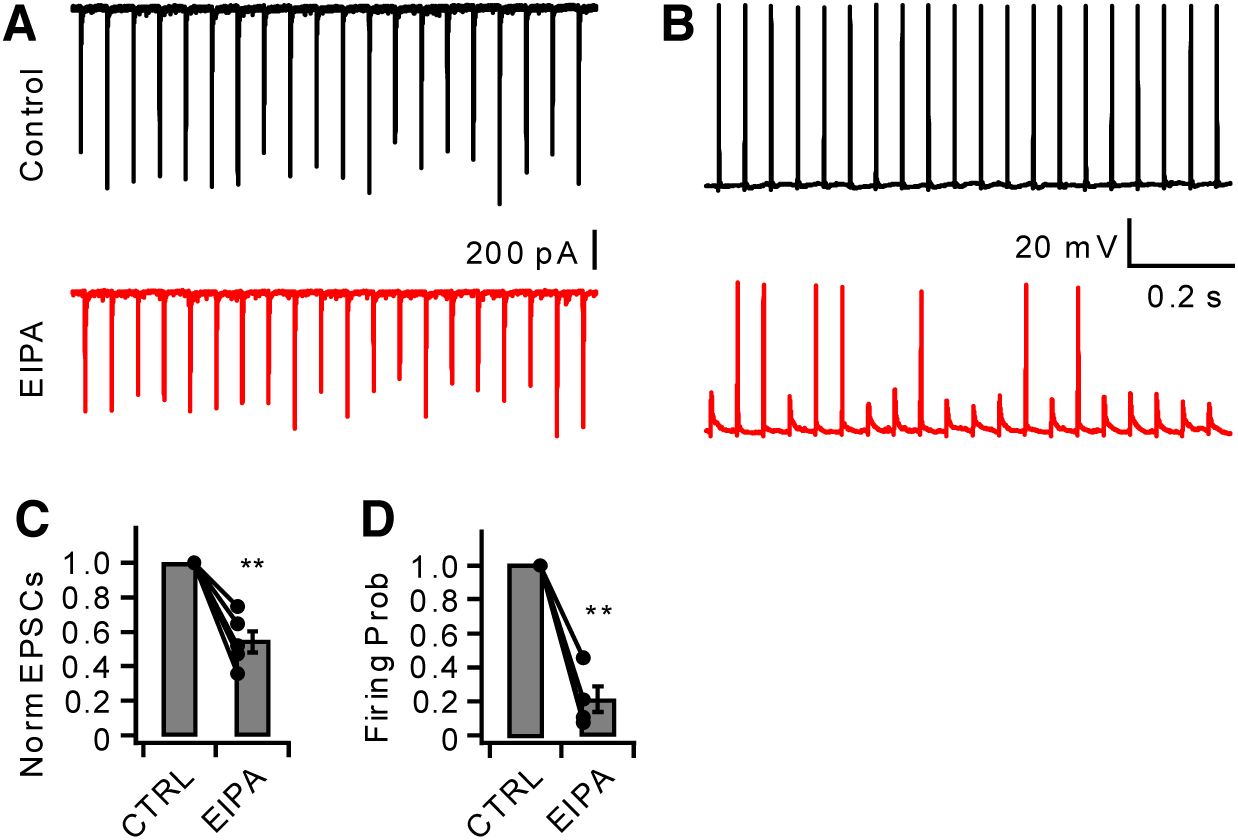
NHE activity is required for reliable synaptic signaling. (A) Example traces of EPSC recordings before and 10 min after incubation of 100 µM EIPA upon 20 Hz presynaptic stimulation. (B) Example traces of postsynaptic action potential recordings before and 10 min after applying EIPA (100 µM). (C) EIPA decreased EPSC amplitude. (D) EIPA reduced reliability of postsynaptic firing. Statistical significance was assessed using a two-tailed paired Student t test. *p < 0.05, **p < 0.01, ***p < 0.001; Error bars, ± SEM.

## Discussion

In this study, we found a substantial accumulation of Na^+^ in the presynaptic terminal during high-frequency signaling. The presynaptic Na^+^ facilitated glutamate uptake into synaptic vesicles without changing the readily releasable pool size or release probability. Our results reveal a mechanism by which action potential-driven Na^+^ influx controls the strength of synaptic transmission by modulating vesicular content. During high-frequency synaptic signaling, when large amounts of glutamate are released, Na^+^ accelerates vesicle replenishment and sustains synaptic transmission, representing a novel cellular mechanism that supports reliable synaptic transmission at high-frequency in the central nervous system.

Glutamate is the principal excitatory neurotransmitter in the brain and is involved in most aspects of brain function (Mayer and Armstrong, 2004). The concentration of glutamate into synaptic vesicles involves plasma membrane excitatory amino acid transporters (EAATs) and vesicular glutamate transporters (VGluTs). EAATs are high-affinity, Na^+^-coupled transporters that recycle glutamate and glutamine from the extracellular space to the cytoplasm; the function of these transporters has been extensively studied (Amara and Fontana, 2002). VGluTs transport glutamate from the cytoplasm into synaptic vesicles for subsequent release by exocytosis, but we still know very little about the basic mechanisms that regulate vesicular glutamate transport, in part because classical biochemical approaches have limitations in revealing the complex ionic basis for the loading of neurotransmitter into vesicles while electrophysiological methods are hard to apply to the study of synaptic organelles (Balmer and Trussell, 2016). Glutamate receptors are not saturated by synaptically released glutamate (Conti and Lisman, 2003; Ishikawa et al., 2002; Sargent et al., 2005); therefore, changes in the amount of glutamate released per synaptic vesicle have the potential to control synaptic strength. Fusion of a single vesicle induces a quantal response, and the size of the quantum varies at most synapses. The variation of miniature EPSC amplitude (quantal size) is determined by vesicular glutamate concentration rather than vesicle volume (Wu et al., 2007), indicating a crucial role of vesicular glutamate uptake in determining synaptic strength. Vesicular transporters drive neurotransmitter accumulation using the energy of the proton electrochemical gradient (Δμ_H+_) produced by the vacuolar H^+^-ATPase (V-ATPase) (Hnasko and Edwards, 2012). The Δμ_H+_ reflects both electrical potential (ΔΨ) and chemical concentration gradient (ΔpH). Previous studies demonstrated that glutamate uptake into synaptic vesicles by the vesicular glutamate transporter (VGluT) is dependent mostly on ΔΨ rather than ΔpH (Maycox et al., 1988; Tabb et al., 1992). However, glutamate entry acidifies synaptic vesicles and reduces the capacity of V-ATPase to create the ΔΨ required for VGluT activity, thereby stalling the uploading of glutamate. Recent work showed that synaptic vesicles express a Na^+^(K^+^)/H^+^ monovalent cation exchanger (NHE) activity that converts ΔpH into ΔΨ and promotes synaptic vesicle filling with glutamate. Manipulating presynaptic [K^+^] changed the mEPSC amplitude even when the glutamate supply was constant, indicating that the synaptic vesicle NHE regulates glutamate release and synaptic transmission (Goh et al., 2011). Another study showed that K^+^ facilitates glutamate transport by directly acting on VGluT binding sites, and confirmed that the NHE is responsible for Na^+^ effects on vesicular glutamate uptake (Preobraschenski et al., 2014). Indeed, presynaptic Na^+^ is more potent than K^+^ in facilitating glutamate uptake and a small change in presynaptic [Na^+^] affect the mEPSC amplitude even in the presence of normal [K^+^]. Na^+^ influx through plasma membrane HCN channels affects presynaptic Na^+^ concentration, regulates glutamate uptake, and thus controls mEPSC amplitudes (Huang and Trussell, 2014). Here we found that Na^+^ facilitated synaptic transmission by facilitating glutamate uptake into synaptic vesicles without changing the readily releasable pool size or release probability. The isoform of NHE It remains unknown. Recent studies suggest NHE6 is likely the candidate: (1) mass spectrometry experiments showed that NHE6 was recently found in both GABAergic and glutamatergic synaptic vesicles (Gronborg et al., 2010); (2) NHE6 was detected by western blotting on highly purified SVs (Preobraschenski et al., 2014); and (3) NHE6 localized to axonal compartments and was increased in area CA1 of the mouse hippocampus during synaptogenesis (Deane et al., 2013).

Na^+^ channels are expressed in hippocampal mossy fiber boutons (Engel and Jonas, 2005), while the location of Na^+^ channels in the calyx of Held is under debate (Kim et al., 2010; Leao et al., 2005; Sierksma and Borst, 2017), although the Na^+^ currents recorded in calyces with incised axon show comparable amplitude (Huang and Trussell, 2008). We found here that Na^+^ increase at the calyceal terminal was comparable (12% smaller) to that of the axon heminode (Fig. 1). Moreover Na^+^ rise and decay kinetics at these locations were identical, supporting that Na^+^ is expressed on or close to the calyx terminals. Although HCN is prominent in controlling the resting Na^+^ concentration, spikes are much more potent in regulating presynaptic Na^+^ accumulation than HCN channels during activity. Blocking HCN channels, which took over 10 min to reach an equilibrated concentration, reduced the resting Na^+^ concentration by about 5 mM (Huang and Trussell, 2014). The calyx Na^+^ increase upon 1 s of 100 Hz firing (Fig. 1) would be larger than the overall contribution of HCN channels at resting membrane potentials. The calyx recorded in brain slices does not fire spontaneously, however, it fires *in vivo* at frequencies of 71 ± 11 Hz in the absence of sound and up to 352 ± 34 Hz with 80 dB tones (Lorteije et al., 2009). Therefore, the presynaptic [Na^+^] would be substantially higher *in vivo* than that of the slice preparations.

Globular bushy cells fire action potentials reliably and precisely synchronize to sound. High-frequency signals of globular bushy cells are reliably transmitted to the target MNTB neurons through the calyx of Held synapse (von Gersdorff and Borst, 2002). Several cellular mechanisms have been established that are important to support neurotransmission at such high rates, including presynaptic ion channels that enable reliable presynaptic spike waveform and calcium influx; a large readily releasable pool, many release sites, and low release probability that enhance the release reliability; and fast kinetics of postsynaptic AMPA-type glutamate receptors that allow fast and faithful transmission to the postsynaptic MNTB neurons (Borst and Soria van Hoeve, 2012; Taschenberger et al., 2002; Taschenberger and von Gersdorff, 2000; Wu et al., 2009). We found that the cytosolic Na^+^-dependent facilitation of vesicular glutamate uptake contributes to the reliability of synaptic transmission at the calyceal synapse. The reliability of action potential propagation from calyx to MNTB dropped to about 50% after a few spikes (Fig. 6D), which is much lower than that in in vivo conditions (Lorteije et al., 2009; Mc Laughlin et al., 2008). This could be explained by the lower presynaptic [Na^+^] when calyx does not fire spike in slice preparation. In the intact brain, however, the calyx fires spontaneously at high frequencies, which would elevate the presynaptic [Na^+^] to over 50 mM (Fig. 1). The elevated [Na^+^] facilitates vesicular glutamate transport and the bigger quantal size ensures reliable synaptic transmission. Indeed, increasing the presynaptic [Na^+^] to 40 mM in the slice preparation rescued the reliability of high-frequency transmission (Fig. 6C-D, left panel). The strength of synaptic transmission is determined by the readily releasable vesicle pool, release probability, and quantal size. Although the readily releasable vesicle pool and release probability reflect presynaptic properties, it is generally accepted that the changes in quantal size indicate postsynaptic alterations in neurotransmitter receptor interactions. Our results show that during high-frequency spiking activity, intracellular Na^+^ is elevated in the terminals and altered the quantal size, which provides a novel presynaptic mechanism to control synaptic strength through changes in the concentration of transmitter in synaptic vesicles without changing the readily releasable vesicle pool or release probability. Since this presynaptic change of synaptic transmission does not affect paired-pulse ratio, our results suggest caution in interpreting studies which use paired-pulse ratio to determine pre- or postsynaptic changes. This activity-dependent modulation of vesicular content provides a novel mechanism of synaptic plasticity, and these findings have potentially universal implications for the regulation of synaptic efficacy in the central nervous system, especially for neurons that fire high-frequency spikes.

## Acknowledgements

We thank Drs. Laurence Trussell, Jeffery Tasker, Laura Schrader, and Youad Darwish for critical reading and comments on the manuscript. Financial supported was provided by U.S. National Institutes of Health grant R01DC016324 to H. Huang.

## Author Contributions

D.L. Y.Z. and H.H. designed, performed, analyzed experiments, and wrote the manuscript. H.H. conceived and supervised the research project.

## Competing Financial Interests

The authors declare no competing financial interests.

